# Growth inhibition by amino acids in *Saccharomyces cerevisiae*

**DOI:** 10.1101/222224

**Authors:** Stephanie J. Ruiz, Joury S. van ’t Klooster, Frans Bianchi, Bert Poolman

## Abstract

Amino acids are essential metabolites but can also be toxic when present at high levels intracellularly. Substrate-induced down-regulation of amino acid transporters in *Saccharomyces cerevisiae* is thought to be a mechanism to avoid this toxicity. It has been shown that unregulated uptake by the general amino acid permease Gap1 causes cells to become sensitive to amino acids. Here, we show that overexpression of eight other amino acid transporters (Agp1, Bap2, Can1, Dip5, Gnp1, Lyp1, Put4 or Tat2) also induces a growth defect when specific single amino acids are present at concentrations of 0.5–5 mM. We can now state that all proteinogenic amino acids, as well as the important metabolite ornithine, are growth inhibitory to *S. cerevisiae* when transported into the cell at high enough levels. Measurements of initial transport rates and cytosolic pH show that toxicity is due to amino acid accumulation and not to the influx of co-transported protons. The amino acid sensitivity phenotype is a useful tool that reports on the *in vivo* activity of transporters and has allowed us to identify new transporter-specific substrates.

## Introduction

As well as being the building blocks for proteins, amino acids provide raw materials for energy generation, nitrogen metabolism, and the biosynthesis of structural, signaling or defensive compounds (Bender 2012). Although they are essential metabolites, it has been known for over half a century that the addition of excess amino acids to both prokaryotic and eukaryotic cell cultures can cause growth inhibition and/or cell death (Rowley 1953a, 1953b; Johnson and Vishniac 1970; Jensen *et al.* 1974; Englesberg *et al.* 1976; Sumrada and Cooper 1976; Nishiuch *et al.* 1976; Miles *et al.* 1976). The dependence of toxicity on transport activity indicate that the effect is exerted intracellularly (Kaur and Bachhawat 2007; Watanabe *et al.* 2014). In humans, several inherited metabolic diseases are caused by elevated levels of amino acids and/or closely related metabolites, with perhaps the most well known being phenylketonuria (Blau *et al.* 2010; Saudubray *et al.* 2016).

Some amino acids are known to cause toxicity via specific mechanisms. Valine and phenylalanine inhibit the growth of bacterial cells by repressing enzymes involved in synthesis of isoleucine and tyrosine, respectively (Leavitt and Umbarger 1962; Bhatnagar *et al.* 1989). Histidine toxicity in yeast cells is caused by a reduction in copper availability *in vivo* (Watanabe *et al.* 2014). There is also evidence for a more general mechanism involving the target of rapamycin complex 1 (TORC1), a master controller of cell metabolism that is conserved among eukaryotes and responds to various nutrient stimuli including amino acids (Duan *et al.* 2015; Powis and De Virgilio 2016; Zheng *et al.* 2016). Deregulation of TORC1 signaling is linked to many human diseases (Laplante and Sabatini 2012), and it has been recently shown that in some cancer cell lines this deregulation is caused by abnormal amino acid transport (Park *et al.* 2016; Cormerais *et al.* 2016; Krall *et al.* 2016). Phenylalanine sensitivity in a mammalian cell line, and rescue by valine, has been shown to involve mTORC1 (Sanayama *et al.* 2014). All amino acids except glutamine are growth inhibitory to the plant *Nicotiana silvestris*, and glutamine itself can rescue some (but not all) of the inhibition caused by other amino acids (Bonner *et al.* 1992, 1996; Bonner and Jensen 1997). Glutamine is now known to be an important activator of TORC1 (Durán *et al.* 2012; Stracka *et al.* 2014; Jewell *et al.* 2015).

Wild-type yeast cells are able to synthesize all amino acids *de novo* but can also scavenge them from the environment using a host of broad- and narrow-range transporters located in the plasma membrane (Ljungdahl and Daignan-Fornier 2012, Gournas *et al.* 2016). *Saccharomyces cerevisiae* amino acid permeases have been studied extensively not only in terms of their transport activity but also their regulation. Many are transcriptionally up- or down-regulated by three interconnected pathways which respond to the availability of amino acids and other nitrogen sources: nitrogen catabolite repression (NCR), general amino acid control (GAAC), and SPS-signaling (Magasanik and Kaiser 2002; Hinnebusch 2005; Ljungdahl 2009; Ljungdahl and Daignan-Fornier 2012). Some transporters are also post-translationally regulated in response to the external concentration of their substrates *e.g.* Can1 (Arg), Dip5 (Glu), Gap1 (various), Lyp1 (Lys), Mup1 (Met), and Tat2 (Trp) (Lin *et al.* 2008; Nikko and Pelham 2009; Hatakeyama *et al.* 2010; Keener and Babst, 2013; Ghaddar *et al.* 2014b). In this pathway, substrate binding triggers transporter ubiquitination and subsequent removal of the proteins from the plasma membrane via endocytosis (MacGurn *et al.* 2012). This causes a decrease in transport activity and thus limits the accumulation of certain substrates. Similar regulation has been observed for Ptr2, which imports di- and tri-peptides that can be broken down into free amino acids inside the cell (Melnykov 2016). It has been suggested that this is a mechanism to avoid amino acid-induced toxicity (Risinger *et al.* 2006; Melnykov 2016).

Risinger *et al.* (2006) showed that *S. cerevisiae* cells expressing Gap1^K9R,K16R^, a ubiquitination- and endocytosis-deficient mutant that is constitutively localized to the plasma membrane, experience severe growth defects when individual amino acids are added to the medium at high concentrations (3 mM). The same effect, with varying levels of severity, was triggered by the metabolite citrulline as well as all proteinogenic amino acids except alanine. We decided to investigate the amino acid sensitivity of strains constitutively overexpressing eight different narrow- and broad-range amino acid transporters: Agp1, Bap2, Can1, Dip5, Gnp1, Lyp1, Put4 or Tat2 (Table 1).

**Table 1.** Transporters used in this study and their reported substrates. Values in brackets indicate published Michaelis constants (*K_m_*).

| Name | Transport substrates | Reference(s)* |
| --- | --- | --- |
| Agp1 | His, Asp, Glu, Ser, Thr, Asn (0.29 mM), Gln (0.79 mM), Cys, Gly, Pro, Ala, Val, Ile (0.6 mM), Leu (0.16 mM), Met, Phe (0.6 mM), Tyr, Trp | Schreve *et al.* 1998; Iraqui *et al.* 1999; Düring-Olsen *et al.* 1999; Andréasson *et al.* 2004; Sáenz *et al.* 2014 |
| Bap2 | Cys, Ala, Val, Ile, Leu (37 μM), Met, Phe, Tyr, Trp | Grauslund *et al.* 1995; Schreve and Garrett 1997; Sáenz *et al.* 2014; Usami *et al.* 2014 |
| Can1 | His, Arg (10–20 μM), Lys (150–250 μM), Orn | Grenson *et al.* 1966; Ghaddar *et al.* 2014a |
| Dip5 | Glu (48 μM), Asp (56 μM), Ser, Asn, Gln, Gly, Ala | Regenberg *et al.* 1998 |
| Gnp1 | Ser, Thr, Asn, Gln (0.59 mM), Cys, Pro, Leu, Met | Zhu *et al.* 1996; Düring-Olsen *et al.* 1999; Andréasson *et al.* 2004 |
| Lyp1 | Lys (10–25 μM), Met | Grenson 1966; Sychrová and Chevallier 1993; Sychrová *et al.* 1993; Ghaddar *et al.* 2014a |
| Put4 | Gly, Pro, Ala | Lasko and Brandriss 1981; Jauniaux *et al.* 1987; Vandenbol *et al.* 1989; Andréasson *et al.* 2004 |
| Tat2 | Gly, Ala, Phe, Tyr, Trp, Cys | Schmidt *et al.* 1994; Düring-Olsen *et al.* 1999; Kanda and Abe 2013 |

\* All proteins in this table were studied in Regenberg *et al.* (1999)

## Materials and methods

### Strains and growth conditions

*Escherichia coli* strain MC1061 was used for cloning and plasmid storage. Experiments were performed with *S. cerevisiae* strains BY4709 (*MATa ura3Δ0*) and BY4741 (*MATa hisΔ1 leu2Δ0 met15Δ0 ura3Δ0*) (Brachmann *et al.* 1998). BY4709 was used for growth experiments and transport assays, while BY4741 was used for measurements of the internal pH.

All experiments were done in YNBD, a minimal medium containing 6.9 g/L Yeast Nitrogen Base without amino acids (Formedium^TM^, UK), 20 g/L D-glucose (Sigma-Aldrich, USA), and 100 mM of potassium phosphate (pH 6.0). Single amino acids or a standard synthetic complete (SC) mixture (Kaiser Drop-out minus uracil, Formedium^TM^) were added for growth assays. For pHluorin experiments a low fluorescence version of YNBD was made using Yeast Nitrogen Base without amino acids and without folic acid and riboflavin (Formedium^TM^) and methionine and histidine were added at 76 mg/L each (509 and 490 μM, respectively). For growth assays shown in Figure 4, the YNBD did not contain potassium phosphate buffer, but the pH was set to 6.0 using HCl/NaOH before sterilization by filtration. All media contain ammonium sulfate (5 g/L) as nitrogen source.

Agar was added at 20 g/L for solid cultures. Liquid cultures (5–15 mL) were inoculated from single colonies on agar plates and incubated in 50 mL CELLreactor^TM^ filter top tubes (Greiner Bio-One, Austria) at 30°C with shaking (~ 200 rpm).

### Plasmids

The plasmids used in this study are listed in Table 2. Vectors for the constitutive expression of transporters are based on pFB022 and pFB023 (Bianchi *et al.* 2016). These are pRS426 derivatives (*URA3*, 2μ) that allow for expression of fluorescently-tagged Lyp1, either full length or truncated, from the constitutive ADH1 promoter (Mumberg *et al.* 1995). The C-terminal tag contains a TEV protease recognition site, followed by the fluorescent protein YPet (Nguyen and Daugherty 2005) and an eight-residue His epitope (for the full sequence, see the Supplementary). Plasmids pSR045–051 are identical to pFB022 except for replacement of *LYP1* with other transporter genes as indicated in Table 1. They were constructed by *in vivo* homologous recombination using crude PCR products. The plasmid backbone was amplified from pFB022 (Bianchi *et al.* 2016), using primers binding immediately upstream and downstream of the *LYP1* coding region. The reaction was then treated with *Dpn*I (as per the manufacturer’s instructions) to remove the template DNA. Each target gene was PCR amplified from *S. cerevisiae* BY4742 chromosomal DNA, using forward and reverse primers that added approximately 25 bp of sequence homologous to the plasmid backbone (Table 3). The backbone and insert were simultaneously transformed into BY4709 and positive transformants screened by growth on uracil dropout media, colony PCR and fluorescence microscopy. Fusion genes were confirmed by sequencing of the entire open reading frame. A single basepair mutation (T to C) was observed at the end of the ADH1 promoter, but this did not appear to affect expression.

**Table 2.**
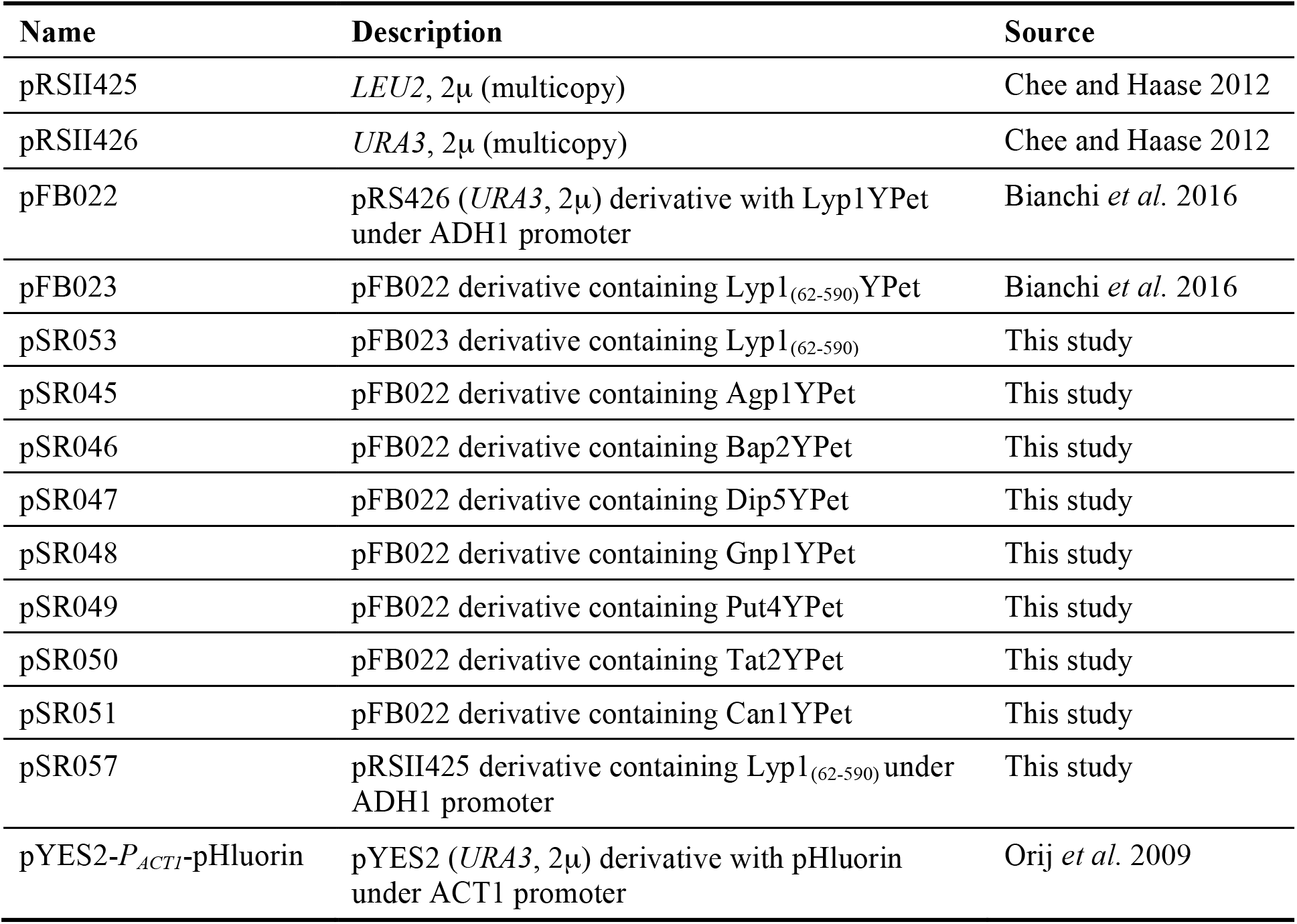
Plasmids used in this study

| Name | Description | Source |
| --- | --- | --- |
| pRSII425 | *LEU2*, 2μ (multicopy) | Chee and Haase 2012 |
| pRSII426 | *URA3*, 2μ (multicopy) | Chee and Haase 2012 |
| pFB022 | pRS426 (*URA3*, 2μ) derivative with Lyp1YPet under ADH1 promoter | Bianchi *et al.* 2016 |
| pFB023 | pFB022 derivative containing Lyp1<sub>(62-590)</sub>YPet | Bianchi *et al.* 2016 |
| pSR053 | pFB023 derivative containing Lyp1<sub>(62-590)</sub> | This study |
| pSR045 | pFB022 derivative containing Agp1YPet | This study |
| pSR046 | pFB022 derivative containing Bap2YPet | This study |
| pSR047 | pFB022 derivative containing Dip5YPet | This study |
| pSR048 | pFB022 derivative containing Gnp1YPet | This study |
| pSR049 | pFB022 derivative containing Put4YPet | This study |
| pSR050 | pFB022 derivative containing Tat2YPet | This study |
| pSR051 | pFB022 derivative containing Can1YPet | This study |
| pSR057 | pRSII425 derivative containing Lyp1<sub>(62-590)</sub> under ADH1 promoter | This study |
| pYES2-*P<sub>ACT1</sub>*-pHluorin | pYES2 (*URA3*, 2μ) derivative with pHluorin under ACT1 promoter | Orij *et al.* 2009 |

**Table 3.**
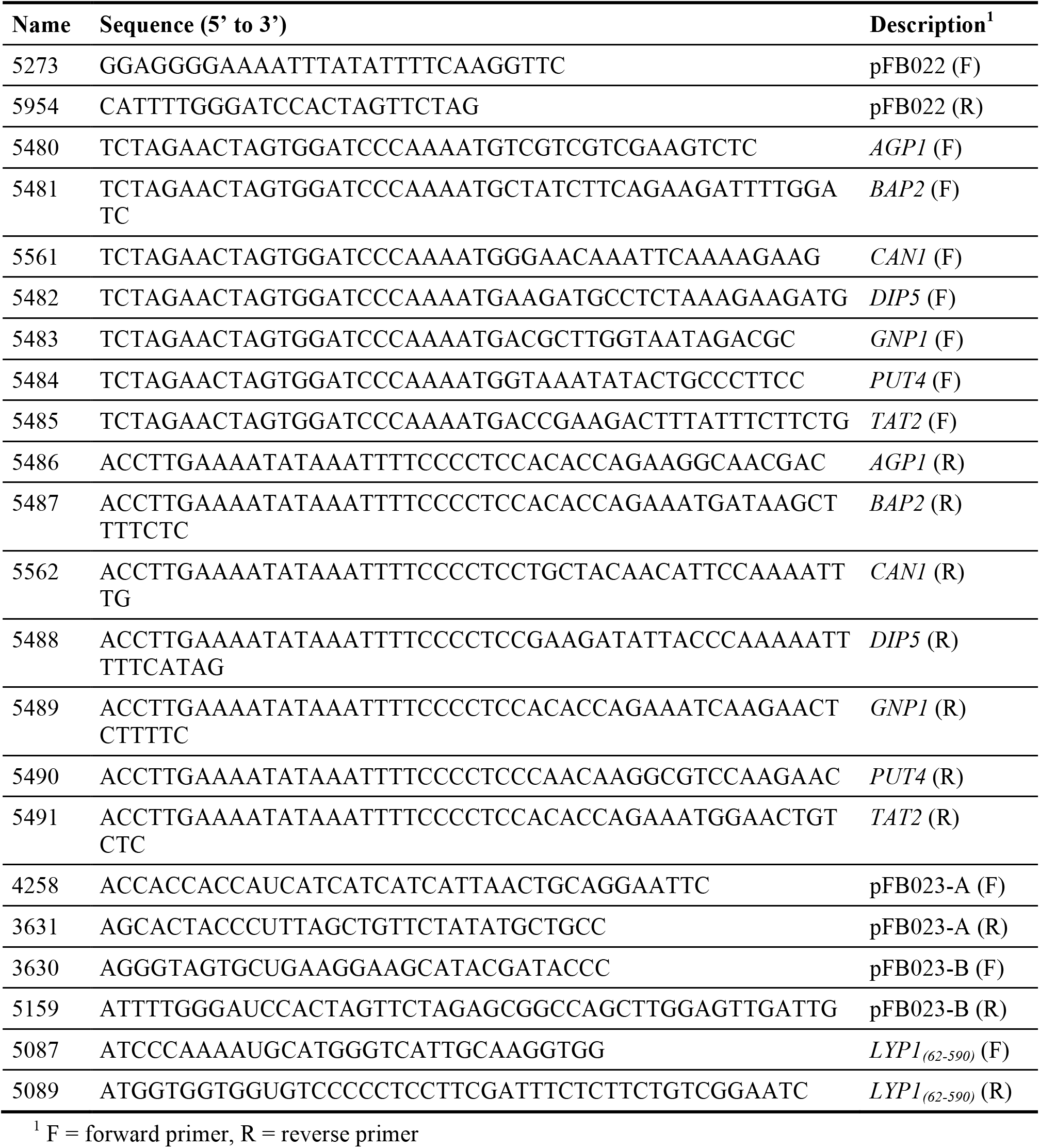
Oligonucleotide primers used in PCR amplification

Ratiometric pHluorin (MiesenbÖck *et al.* 1998) was expressed from pYES2-P_ACT1_-pHluorin (Orij *et al.* 2009). For these experiments a truncated Lyp1 with no fluorescent tag was used (Bianchi *et al.* 2016). First, the C-terminal YPet tag was removed from pFB023 using uracil excision-based cloning (Bitinaite and Nichols 2009). Primer pairs 4158/3631, 3630/5159 and 5087/5089 (Table 3) and the PfuX7 polymerase (Nørholm 2010) were used to amplify pFB023 in three fragments excluding the YPet coding region. The crude PCR products were treated with USER™ enzyme (New England Biolabs, USA) as per the manufacturer’s instructions and transformed into *E. coli* MC1061 for *in vivo* assembly. After the resulting plasmid (pSR053) was confirmed by sequencing of the entire coding region, the area containing *LYP1_(62-590)_* and the flanking promoter and terminator sequences were sub cloned into pRSII425 using *SacI/KpnI* digestion to yield plasmid pSR057. The expressed protein (from N-to C-terminus) includes a starting Met, residues 62-590 of Lyp1, a short three-residue glycine linker, and an eight-residue His epitope.

### Microscopy

Live cell imaging was performed on a LSM 710 commercial laser scanning confocal microscope (Zeiss, Germany) equipped with a C-Apochromat 40 x/1.2 NA objective. Cells were immobilized between a glass slide and coverslip. Images were obtained with the focal plane positioned at the mid-section of the cells. For fluorescence images, samples were excited with a blue argon ion laser (488 nm) and emission collected at 493–797 nm.

### Growth assays

Growth assays were performed using 120 μL liquid cultures in CELLSTAR® 96-well flat-bottom microplates (Greiner Bio-One). Each plate was covered with a Breathe-Easy® sealing membrane (Sigma-Aldrich), as well as the provided lid, and incubated in a 30°C room at 400 rpm on an Excella™ E1 benchtop open-air shaker (Eppendorf, Germany). OD_600_ measurements were made in a PowerWave 340 spectrophotometer (BioTek, USA) without the microplate lid but with the (optically clear) membrane.

To prepare inocula for microplate experiments, strains were cultured in YNBD for approximately 24 h, with one round of dilution, to an OD_600_ of 0.3–0.7. Each culture was diluted in fresh media to an OD_600_ of 0.1 and 60 μL aliquots were added to microplate wells containing 60 μL of media with or without either single amino acids or the SC mixture. The final concentrations are given in Table 4 and Figures 2 and 4. All measurements were done in biological triplicate (three independent inoculations made on different days from different precultures).

Raw OD_600_ values were background corrected by subtracting the average value for all blank (media only) wells from the same plate (n = 12–24). This observed value (ODobs) was then corrected for non-linearity at higher cell densities using the polynomial equation: OD_cor_ = 0.319 × ODobs^3^ + 0.089 × ODobs^2^ + 0.959 × ODobs (Figure S1); this correction method was previously discussed in Warringer and Blomberg (2003). OD_cor_ values were then normalized within each strain and replicate. Three independent experiments were performed but due to technical error some results had to be discarded. For this reason, n = 2 for Bap2 1 mM Gly/His/Ile/Leu/Lys/Met/Orn/Phe, and also for all Tat2 1 mM and 0.5 mM conditions.

**Table 4.** Concentration (mM) of components in synthetic complete (SC) mixture, when used at 1x.

| **Amino acids** |  |
| --- | --- |
| Alanine (Ala) | 0.853 |
| Arginine (Arg) | 0.361 |
| Asparagine (Asn) | 0.575 |
| Aspartic acid (Asp) | 0.571 |
| Cysteine (Cys) | 0.627 |
| Glutamine (Gln) | 0.520 |
| Glutamate (Glu) | 0.517 |
| Glycine (Gly) | 1.012 |
| Histidine (His) | 0.490 |
| Isoleucine (Ile) | 0.579 |
| Leucine (Leu) | 2.897 |
| Lysine (Lys) | 0.520 |
| Methionine (Met) | 0.509 |
| Phenylalanine (Phe) | 0.460 |
| Proline (Pro) | 0.460 |
| Serine (Ser) | 0.723 |
| Threonine (Thr) | 0.638 |
| Tryptophan (Trp) | 0.372 |
| Tyrosine (Tyr) | 0.419 |
| Valine (Val) | 0.649 |
| **Other** |  |
| Adenine | 0.133 |
| *myo*-Inositol | 0.422 |
| 4-Aminobenzoic acid | 0.058 |

### Transport assays

*In vivo* transport assays were performed as previously described (Bianchi *et al.* 2016). The radioactive substrates used were L-(^14^C(U))-phenylalanine, L-(^14^C(U))-lysine and L-(methyl-^3^H)-methionine (PerkinElmer, USA) and L-(^14^C(U))-alanine (Amersham Biosciences, UK). Transport was assayed at the following concentrations: 50 μM lysine with and without 100 mM of histidine or ornithine, 500 μM alanine or methionine, and 2.5 mM phenylalanine.

### Measurement of cytosolic pH

Strains were cultured in liquid media to an OD_600_ of 0.3–0.7, diluted to an OD_600_ of 0.2 in pre-warmed media, and then immediately transferred to the spectrophotometer sample holder (also pre-warmed). After 15 minutes, 10 μL of either distilled water or 100 mM lysine was added (final concentration 500 μM) and the measurement continued for another 3 h.

Fluorescence measurements were made using a JASCO FP-8300 fluorescence spectrophotometer with the following settings: sensitivity, high; response, 0.1 s; excitation/emission bandwidths, 5 nm; emission, 508 nm; excitation, 355–495 nm in 5 nm steps. All samples were 2 mL in 4.5 mL plastic cuvettes (catalogue number 1961, Kartell Labware, Italy) with a magnetic bar for stirring. The temperature of the sample holder was maintained at 20°C for calibration measurements and 30°C for experiments with growing cells.

To prepare calibration samples, strains were cultured as above. Cells were then washed twice and resuspended in 5 volumes of ice-cold PBS (10 mM Na2HPO4, 137 mM NaCl, 2.7 mM KCl, 1.8 mM KH_2_PO_4_, pH 7.35). Digitonin was added to a final concentration of 0.02% w/v (from a 2% w/v stock in PBS) and the samples incubated for 1 h at 30°C with mixing. Cells were then washed twice with 1 mL of ice-cold PBS, resuspended to 0.25 mg/mL with the same (1 μL added per mg wet weight of cell pellet) and kept on ice. 10 μL of cells was added to 2 mL of room-temperature buffer (either 100 mM potassium phosphate or PBS) and a fluorescence measurement made once per minute for ten minutes.

We found that background subtraction was very important, especially for long-term measurements with growing cells. Strains carrying pHluorin/pRSII425 or pHluorin/Lyp1 were compared to strains carrying pRSII426/pRSII425 and pRSII426/Lyp1, respectively. To generate the calibration curve (Figure S2), the median emission intensities at 395 and 475 nm excitation (I395 and I475) were background subtracted and the ratio (R395/475) plotted against the pH. For measurements with growing cells, the I395 and I475 values from each strain and timepoint were background subtracted using individual values from the corresponding strain and timepoint.

## Results and discussion

### All standard proteinogenic amino acids, as well as ornithine and citrulline, inhibit growth of *S. cerevisiae*

Eight different *S. cerevisiae* amino acid transporters were overexpressed by introducing their genes on multicopy plasmids under control of the constitutive ADH1 promoter (Figure 1). A C-terminal tag containing the fluorescent protein YPet was added to allow visualization of the transporters in the cell. Confocal fluorescence microscopy confirmed that all eight transporters localize primarily to the cell periphery (Figure 1A), indicating that our tagged proteins are delivered to the plasma membrane. Some fluorescent signal is seen in internal membranes and the vacuole, which is not uncommon for plasma membrane transporters given that their normal life cycle involves being trafficked between these different locations (O’Donnell *et al.* 2010; MacGurn *et al.* 2012). We observed substantial variation in protein expression between cells from the same culture, with a significant proportion showing little or no fluorescence (Figure S3). We believe that this reflects a sub-population of cells that have lost the expression vector. For the high copy pRS plasmids it is known that plasmid-free cells make up 20–30% of the population, even in selective media (Christianson *et al.* 1992). Expression of particular proteins can increase this population significantly, with reported values of up to 50% (Ro *et al.* 2008). Using flow cytometry, we found that the fraction of cells in the fluorescent population (30—65%) and the median fluorescence signal from individual cells in this population (1.5—5.6 arbitrary units) was reasonably stable between independent cultures of each strain (Figure 1B and C). In minimal media without amino acids, growth of the overexpression strains over 24 h was up to 40% lower than that of BY4709 containing the empty plasmid pRSII426 (Figure 1D). This is likely due to both the plasmid instability discussed earlier (different populations of plasmid-free cells translates to a difference in the starting cell density of each culture), as well as general effects caused by overexpression (Österberg *et al.* 2006; White *et al.* 2007). These variations within and between strains do not affect the conclusions of this paper as we only make comparisons of the same strains grown in different conditions.

**Figure 1.**
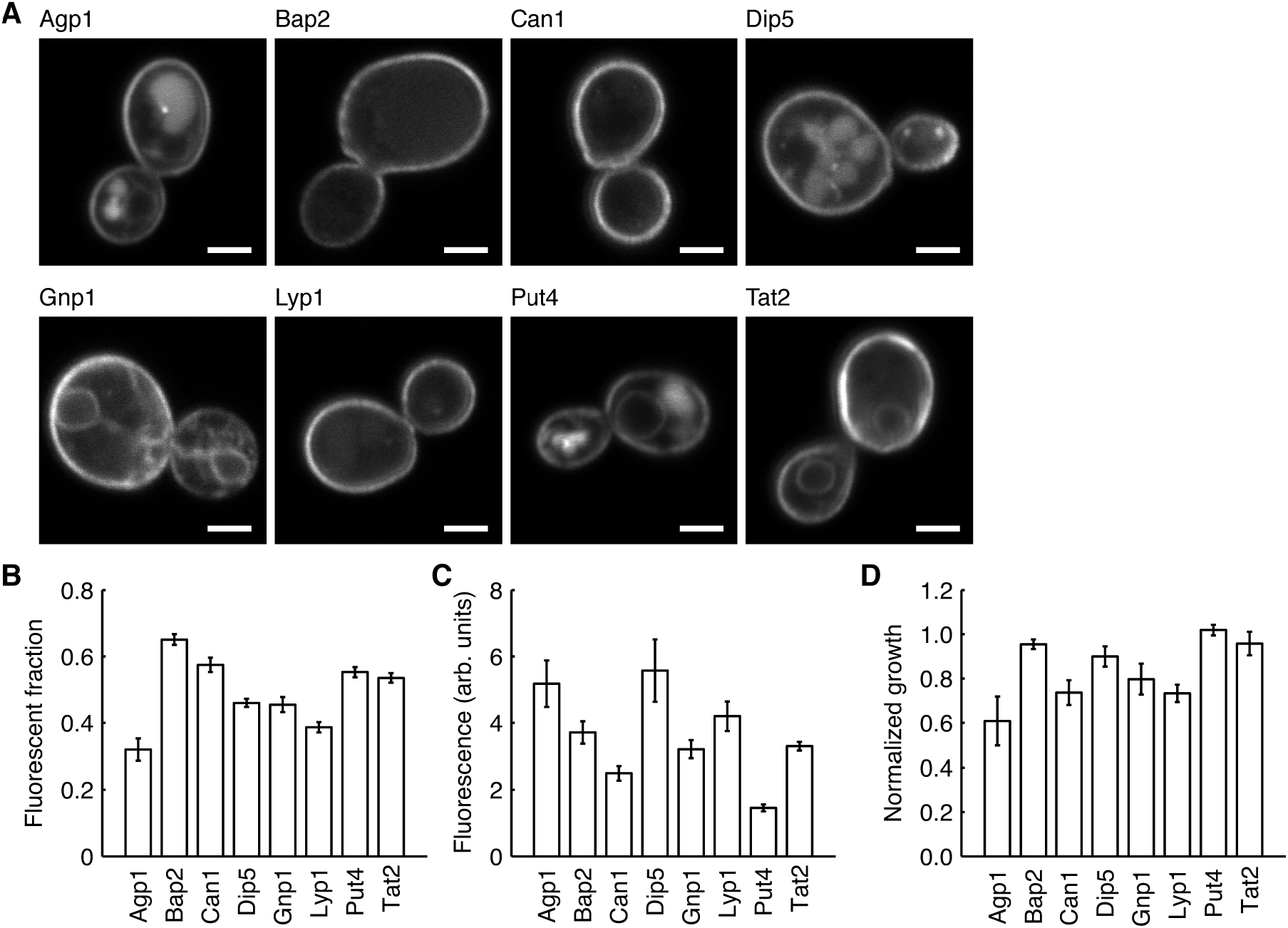
Overexpression of amino acid permeases in BY4709. All proteins contained a C-terminal YPet tag and were expressed from a multicopy plasmid under control of the constitutive ADH1 promoter. (A) Fluorescence confocal microscopy images showing that the transporters localize mainly to the cell periphery. Scale bars are 2 μM. (B, C) Data from flow cytometry analysis showing the estimated fraction of fluorescent cells in each culture (B), and the median fluorescence signal of cells in this sub-population (C). (D) Growth of overexpression strains in minimal media. Normalized growth refers to cell density after 24 h, normalized to BY4709 carrying the empty plasmid pRSII426. Values shown are the mean ± standard deviation (n = 5 for B and C, n = 7 for D).

We compared the growth of each strain in minimal medium with or without the addition of either single amino acids or a synthetic complete (SC) mixture (Figure 2). It should be noted that BY4709 contains all the endogenous *S. cerevisiae* amino acid transporters and is in the yeast S288C background. We therefore expect that at the start of the experiment all strains, which were precultured in minimal medium containing ammonium as the sole nitrogen source, would contain significant levels of active Gap1 (Courchesne and Magasanik 1983; Chen and Kaiser 2002). For the control strain, the SC mixture increased growth and only cysteine or histidine at 5 mM was inhibitory. For all overexpression strains except Put4, at least two different individual amino acids (other than cysteine or histidine) inhibited growth by more than 50%. We observed no citrulline-mediated growth inhibition, consistent with the fact that Gap1 is the only citrulline transporter (Grenson *et al.* 1970). Risinger *et al.* (2006) observed no alanine-mediated toxicity and only a 20% reduction in cell growth caused by phenylalanine (all other amino acids caused a ≥ 65% reduction in growth). Here, we show that 5 mM of phenylalanine reduces the growth of Agp1 and Gnp1 strains by 70%, while 5 mM of alanine reduces the growth of Agp1 strains by 30%. Ornithine, a basic amino acid and important metabolite, inhibited the growth of strains overexpressing Can1 and Lyp1.

**Figure 2.**
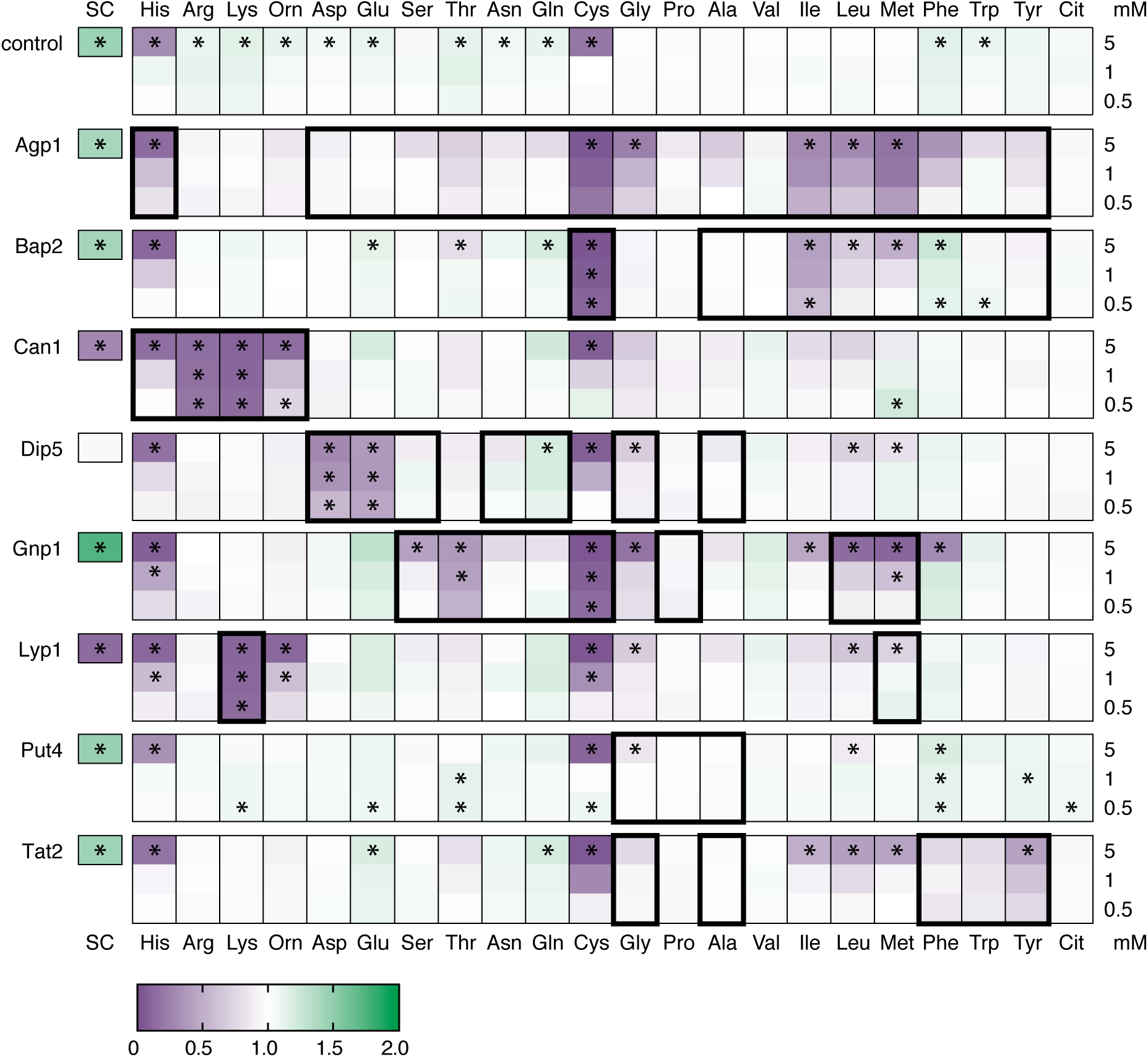
The effect of amino acids on the growth of strains overexpressing amino acid transporters. BY4709 carrying the empty plasmid pRSII426 was used as a control. All proteins were expressed from the constitutive ADH1 promoter with a C-terminal YPet tag. Heat maps show cell density (OD_600_) after 24 h, normalized such that the same strain in YNBD minimal media without amino acids = 1. Raw values are given in Figure S4. The proteinogenic amino acids are represented by their three letter codes (see Table 4). Orn, ornithine; Cit, citrulline; SC, synthetic complete media minus uracil. Amino acids were added at final concentrations of 0.5, 1, or 5 mM. Thick black boxes indicate reported substrates for each transporter (for references see Table 1). Asterisks indicate *p* < 0.05 when compared to growth in YNBD (*t*-test).

We were surprised that proline or valine did not cause higher levels of growth inhibition. Both of these amino acids caused a 90% reduction in growth when supplied at 3 mM to an *S. cerevisiae* strain expressing a endocytosis-resistant Gap1 mutant (Risinger *et al.* 2006). Agp1, Gnp1 and Put4 are all proline transporters, with Put4 estimated to have an approximately 500-fold lower *K*_m_ for proline than Gap1 (Lasko and Brandriss 1981; Andréasson *et al.* 2004). Valine is known to be a substrate for both Agp1 and Bap2, with transport in sub-mM concentrations occurring at rates equal or higher to that of other substrates for which we did observe growth inhibition (e.g. Ile, Leu) (Grauslund *et al.* 1995; Iraqui *et al.* 1999; Regenberg *et al.* 1999). Sensitivity to other amino acids and increases in whole cell uptake (Figure 2 and 3A) indicate that Agp1, Bap2 and Gnp1 are all present in these strains. The largest effect caused by proline or valine, however, was an approximately 10% reduction in the growth of Agp1 strains. It is possible that this difference is due to variations in substrate-regulated trafficking. Although the use of a constitutive promoter minimizes transcriptional regulation, the levels of transporter at the plasma membrane and thus the rate of substrate transport and accumulation is subject to post-translational regulation via intracellular trafficking (Lin *et al.* 2008; Nikko and Pelham 2009; Hatakeyama *et al.* 2010; Ghaddar *et al.* 2014b). A series of elegant experiments using Can1 and Gap1 mutants showed that this process requires ligand binding but not transport, supporting the hypothesis that substrate binding induces conformational changes that promote endocytosis (Ghaddar *et al.* 2014b). The same study demonstrated that different substrates are more or less effective at triggering endocytosis of the same transporter, and it follows that the same amino acid could be more or less effective at triggering endocytosis of different transporters.

**Figure 3.**
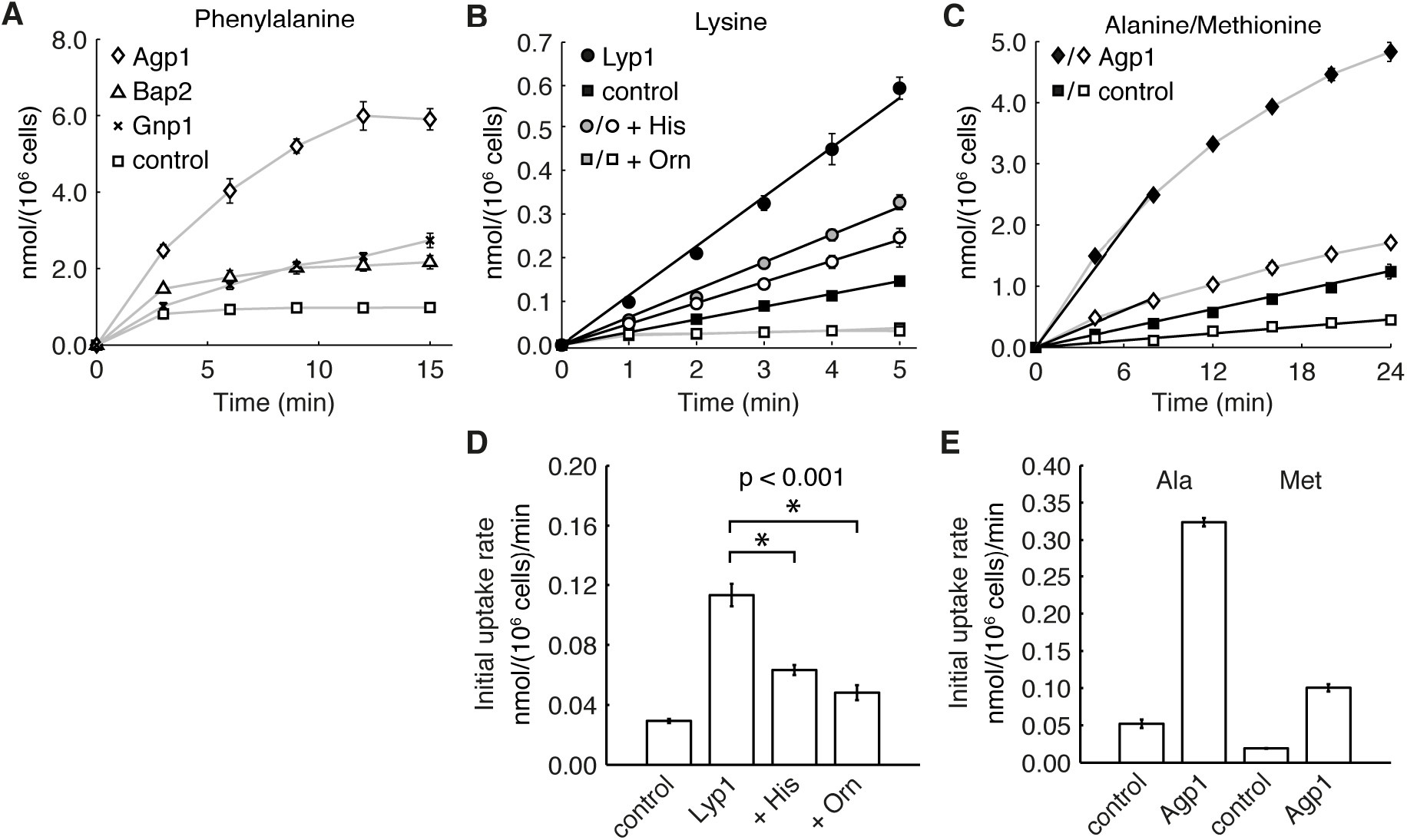
Amino acid uptake by *S. cerevisiae* cells. Transporters were expressed from the constitutive ADH1 promoter with a C-terminal YPet tag. BY4709 carrying the empty plasmid pRSII426 was used as a control. (A) Uptake of phenylalanine (2.5 mM). (B) Uptake of lysine (50 μM) in the absence or presence of histidine or ornithine (100 mM each). (C) Uptake of either alanine (500 μM) or methionine (500 μM). The straight black lines in (B) and (C) represent the calculated initial uptake rates, which are shown in (D) and (E), respectively. Values are the mean of biological triplicates. Error bars represent standard deviation and in some cases are obscured by the data point. Rates in (D) were compared using a *t*-test.

### Amino acid sensitivity reports on transporter activity and substrate specificity

We expected that the pattern of amino acid sensitivity for each strain would match the known substrate specificity (Table 1) of the overexpressed transporter and for the most part this was true (Figure 2). As discussed in the previous section, some known substrates did not cause substantial growth inhibition. Even more surprising was that many strains were sensitive to amino acids not predicted to be transport substrates. While it is possible that these results are affected by altered expression levels of endogenous systems, we tentatively conclude that these transporters are able to transport a much broader range of substrates than previously described, but with a high Michaelis constant (*K*_m_ > 1 mM). This can be rationalized by structural and mutational studies, which indicate that the residues interacting with the α-amino and α-carboxyl groups of the substrate are conserved (Kanda and Abe 2013; Usami *et al.* 2014; Ghaddar *et al.* 2014a). The reason why this broader specificity has not been seen before could be simply because previous studies have tested a limited range of substrates and/or concentrations. The screening study published by Regenberg *et al.* (1999), for example, only assayed transport at 100 or 250 μM of substrate. These lower-affinity transport activities would be important to consider in a laboratory setting where synthetic media often contains amino acids at (sub-)mM concentrations.

Some of these novel substrate specificities (the transport of Phe by Gnp1 and of His and Orn by Lyp1) were further investigated by monitoring the transport of radioactive substrates into whole cells (Figure 3). Again it should be noted that some endogenous transporters, including Gap1, are expected to be present under these conditions but rapidly endocytosed in response to amino acid addition. For this reason we only measured initial rates of transport. Overexpression of Gnp1 indeed increased the uptake of phenylalanine into whole cells (Figure 3A), supporting its identification as a phenylalanine transporter. Lysine uptake was 4-fold faster in cells overexpressing Lyp1 (Figure 3B and D), indicating that the YPet-tagged Lyp1 is active. When His or Orn was present at 100 mM (2000-fold excess) Lys transport by Lyp1 cells was reduced by 56% and 43%, respectively. At the same concentrations, Lys transport by the control strain was reduced to background levels (Figure 3B). These results, in conjunction with the growth inhibition, are consistent with Lyp1 transporting both His and Orn. They also suggest that Lyp1 has a lower affinity for these substrates than Can1, which has a reported inhibition constant (*K*_i_) of 3 mM for both His and Orn (Grenson *et al.* 1966), or Gap1. It has previously been reported that Lyp1 does not transport Orn or His, but this could be explained by a lower affinity and the fact that competition for Lys transport by Lyp1 was assayed with only a ten-fold excess in comparison to the 1000-fold excess used in the competition for Arg transport by Can1 (Grenson *et al.* 1966; Grenson 1966). Subsequent studies using Lyp1 overexpression strains tested for His transport using concentrations of 50–100 μM, which is likely to be far below the *K*_m_ (Sychrová and Chevallier 1993; Regenberg *et al.* 1999).

In further experiments using lower concentrations, Lyp1 strains were inhibited (≥ 50% reduction in growth) by 16 μM lysine and Can1 strains by 16 μM of arginine or 125 μM of lysine (Figure 4). This is in the range of the measured *K*_m_ values (Table 1) (Ghaddar *et al.* 2014a; Bianchi *et al.* 2016). The growth advantage seen for Lyp1 overexpression strains in the presence of external arginine is due to import of this amino acid, which can be used as a carbon or nitrogen source, by other endogenous transporters. The same effect is observed for the control strain, although it is only apparent during the exponential growth phase (Figure S5) and thus not seen at the 24 h time-point used in Figure 4.

**Figure 4.**
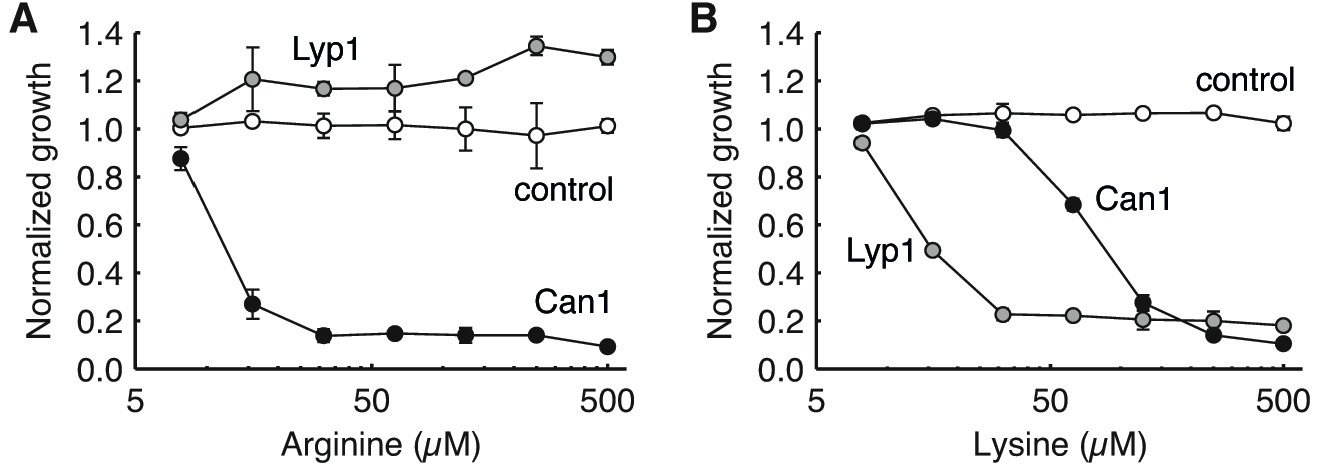
The effect of external arginine (A) and lysine (B) on the growth of strains overexpressing Can1 or Lyp1. BY4709 carrying the empty plasmid pRSII426 was used as a control. Can1 and Lyp1 were expressed from the constitutive ADH1 promoter with a C-terminal YPet tag. Normalized growth refers to cell density after 24 h, normalized to minimal media without amino acids. Values are the mean of biological triplicates. Error bars represent standard deviation and in some cases are obscured by the data point.

### Toxicity is caused by amino acid accumulation, not proton transport

The amino acid transporters studied here, as well as Gap1, are thought to be amino acid:proton symporters. One possible explanation for the toxicity of amino acid transport is that the corresponding influx of protons interferes with cellular homeostasis. This has been previously demonstrated in *E. coli* where excessive, uncontrolled transport of galactosides by the lactose:proton symporter LacY decreases the electrochemical proton gradient across the cell membrane and lowers the intracellular ATP concentration (Wilson *et al.* 1980, 1981). In the case of *S. cerevisiae* amino acid transport, it is possible that this effect would be amplified by the transport of excess amino acids into the vacuole, which is mediated by amino acid:proton antiporters and would thus cause further movement of protons into the cytoplasm (Ohsumi and Anraku 1981; Sato *et al.* 1984).

Risinger *et al.* (2006) argued that proton influx was not the mechanism for Gap1-mediated amino acid toxicity by showing that growth was not inhibited by amino acid mixtures, a condition where the overall transport rate should remain high but individual amino acids would accumulate to lower levels. We did observe toxicity when cells were grown in a mixture of amino acids; in SC media the growth of Can1 and Lyp1 strains was reduced 70% and 85%, respectively, while Dip5 did not have the growth advantage seen for the control and other overexpression strains (Figure 2). However this likely reflects the narrow specificities of these transporters and their relatively low K_*m*_. The other amino acids are presumably not present in high enough concentrations to effectively compete for transport, and thus one or two substrates are still accumulated to high levels. Whole cell transport assays did not show any direct correlation between growth inhibition and initial transport rate (Figure 3), and this suggests that toxicity is not mediated by proton influx. Overexpression of Agp1 increased the transport of both Ala and Met 5- to 6-fold when each amino acid was supplied at 500 μM (Figure 3C and E), with the initial transport of Ala 3-fold higher than that of Met, yet at the same external concentration only Met inhibits growth. 50 μM of Lys is inhibitory to Lyp1 overexpression strains, yet the initial rate of transport is less than half that of the transport of Ala by Agp1 strains (Figure 3D and E). These transport assays also suggest differences in the intracellular concentration at which various amino acids become toxic. Assuming an internal volume of 70 fL per cell (Sherman 2002), we calculated that the amino acid pool of the Agp1 overexpression strain increased by 69 mM of Ala and 24 mM of Met in 24 min, and by 84 mM of Phe in 15 min. Under the same conditions Met and Phe, but not Ala, cause growth inhibition of this strain.

To investigate the effect of increased proton flux more directly, we used a pH-sensitive GFP variant called ratiometric pHluorin (Miesenböck *et al.* 1998; Orij *et al.* 2009) to monitor the cytoplasmic pH (pHc) in growing cells with or without the addition of amino acid to the media (Figure 5A). For these experiments a truncation mutant of Lyp1 was used which is more stably maintained at the plasma membrane after lysine addition (Bianchi *et al.* 2016). Overexpression of this variant also causes lysine sensitivity (data not shown). For control cells or Lyp1 cells without the addition of lysine, the pH_c_ was 7.0–7.2 and stayed fairly constant over three hours. In contrast, the pH_c_ of Lyp1 cells given 500 μM of lysine began to decrease after one hour, and continued to drop steadily over the course of the experiment with a total change of 0.4 pH units. Neither the magnitude of the pH change or the time scale on which it happens is consistent with the hypothesis of rapid proton influx as a causative mechanism. When yeast are exposed to glucose after a period of starvation, the cytoplasmic pH transiently drops to as low as pH 6 and cells are able to recover to normal levels within minutes (Dechant *et al.* 2010; Tarsio *et al.* 2011). Several groups have demonstrated a direct correlation between pHc and growth rate in yeast, but *S. cerevisiae* is still able to grow reliably at pH_c_ between 6.5 and 7 (Orij *et al.* 2012; Dechant *et al.* 2014).

**Figure 5.**
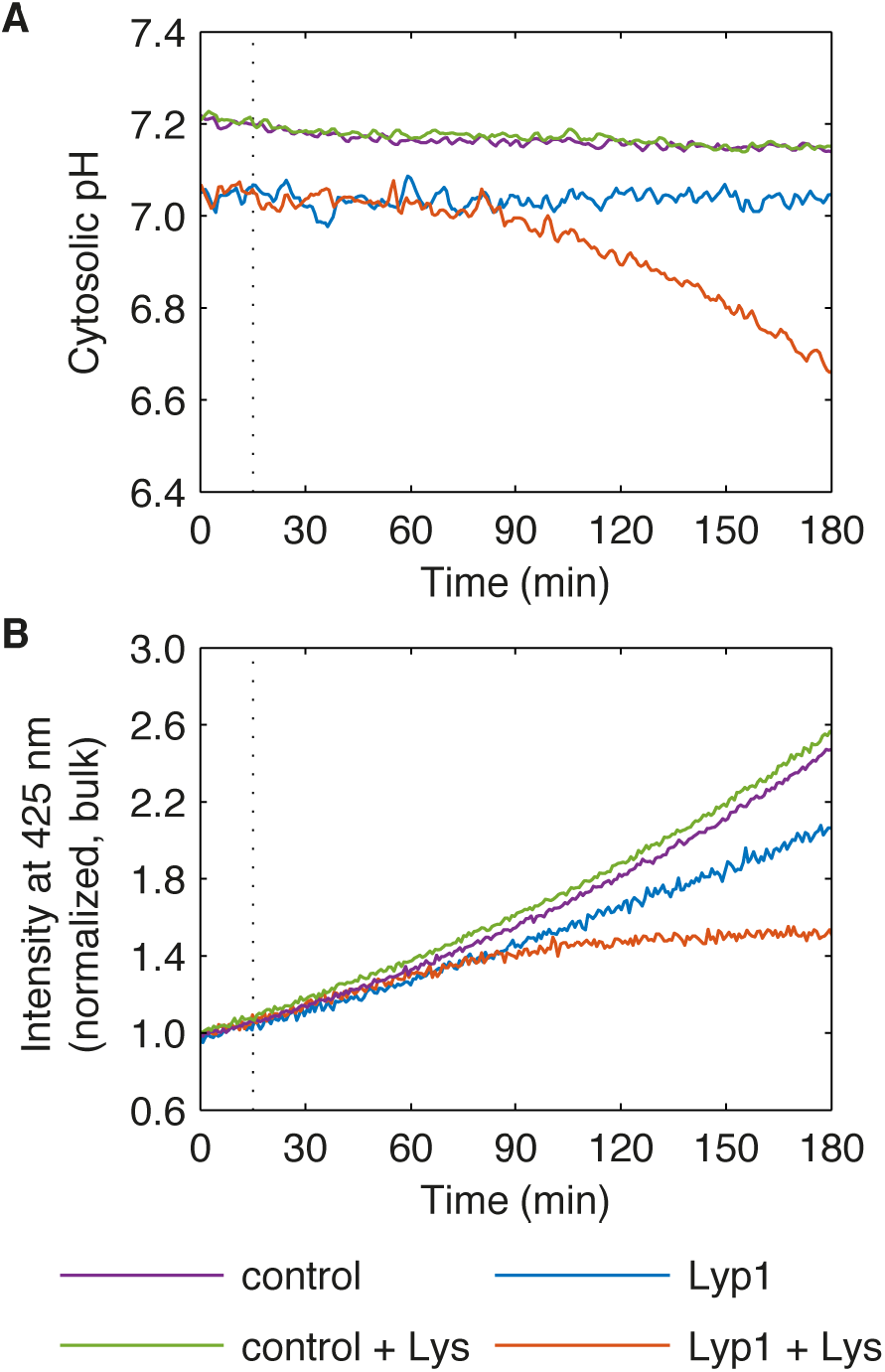
The effect of external lysine on the cytosolic pH of cells overexpressing Lyp1. BY4741 expressing the pH-sensitive ratiometric pHluorin and carrying the empty plasmid pRSII425 was used as a control. The Lyp1 strain constitutively expresses both pHluorin and a truncated, non-fluorescently tagged Lyp1. (A) Cytosolic pH. (B) Normalized bulk intensity at 425 nm excitation/508 nm emission, at which pHluorin is largely insensitive to pH. Values shown are the mean of two independent experiments (individual results are shown in Figure S6). Dotted lines indicate the time at which either distilled water or lysine (final concentration 500 μM) was added.

We were able to obtain more information from the pHluorin experiments by analyzing the fluorescence signal at 425 nm excitation/508 nm emission. Under these conditions pHluorin is insensitive to pH (Miesenböck *et al.* 1998). This means that we can monitor the amount of pHluorin independent of the pH changes. For the samples where pHc remained constant, the bulk fluorescence fits well to an exponential curve (Figure 5B). We believe that this value is reporting the combined rates of protein production plus degradation in dividing cells, with the amount of pHluorin in individual cells staying stable over time. In contrast, the bulk fluorescence of Lyp1 cells after lysine addition begins to plateau at approximately the same time that the cytosolic pH begins to decrease. This suggests that the mechanisms leading to growth inhibition are occurring in the hour after lysine addition, and that the decrease in cytosolic pH is a consequence, rather than a cause.

One hypothesis that would fit our data is that lysine (and also cysteine) toxicity is due to interference with ubiquitination pathways. Protein modification by the attachment of ubiquitin (Ub) is a regulatory mechanism involved in a wide range of essential processes in eukaryotic cells (Finley *et al.* 2012). A key step in these pathways is the transfer of Ub from a Ub-conjugating (E2) enzyme to a target protein. Some human E2~Ubs are able to react with free lysine and cysteine molecules in such a way that Ub is irreversibly transferred to the amino acid (Pickart and Rose 1985; Wenzel *et al.* 2011). Intrinsic reactivity with free lysine has also been observed for the *S. cerevisiae* E2 enzymes Ubc4 and Pex4 (Chris Williams, personal communication). This reaction occurs on the minute timescale, in the cytoplasmic pH range and at concentrations as low as 50 mM. Extrapolation of our transport assay data suggests that our overexpression strains import this amount of lysine within 30–60 minutes, which is in agreement with previous work (Bianchi *et al.* 2016). Overlapping activities mean that individual E2 enzymes are not essential in *S. cerevisiae* but multiple knockouts, for example *Δubc4Δubc5*, are lethal (Stoll *et al.* 2011).

## Conclusions

Our study expands on the work of Risinger *et al.* (2006) who showed that all 20 proteinogenic amino acids, as well as the non-proteinogenic amino acids citrulline and ornithine, can be growth inhibitory to *S. cerevisiae*. We have shown that this effect can be mediated by various amino acid transporters and is not specific to Gap1. We have also demonstrated that amino acid-mediated growth inhibition is not dependent on the initial rate of transport, or triggered by the rapid influx of protons, but is instead caused by the longer-term accumulation of single amino acids.

Amino acid sensitivity is an important phenomenon that should be considered in the design and analysis of studies of amino acid and peptide transport. It is also a useful tool for assessing the *in vivo* activity of transporters as it reports on their levels at the plasma membrane and their transport kinetics for specific substrates. We have used it to develop growth-based screens to confirm the activity of overexpressed wild-type and mutant transporters, including when expressed in *Pichia pastoris* (Bianchi *et al.* 2016). Our results from screening eight different amino acid transporters did vary from what we predicted based on the current literature suggesting (i) a much broader specificity than previously thought, and (ii) transporter/substrate specific variations which may reflect differences in substrate-induced down-regulation.

## Acknowledgements

We thank Dr Gertien J. Smits for providing the plasmid pYES2-P_ACT1_-pHluorin, and Dr Chris Williams for general discussion of our work.

The authors declare that there is no conflict of interest.

## Supplementary

The C-terminal tag, whose full amino acid sequence is shown below, contains a TEV protease recognition site (underlined) followed by the fluorescent protein YPet (in bold) and an eight-residue His epitope (in italics). The residue pairs “GG” and “EL” are a short linker and a cloning artifact, respectively.

<u>GGENLYFQG</u>**SKGEELFTGVVPILVELDGDVNGHKFSVSGEGEGDATYGKLTLKLLCTTGKLP VPWPTLVTTLGYGVQCFARYPDHMKQHDFFKSAMPEGYVQERTIFFKDDGNYKTRAEVKFEG DTLVNRIELKGIDFKEDGNILGHKLEYNYNSHNVYITADKQKNGIKANFKIRHNIEDGGVQL ADHYQQNTPIGDGPVLLPDNHYLSYQSALFKDPNEKRDHMVLLEFLTAAGITEGMNELYK**EL *HHHHHHHH*

**Figure S1.**
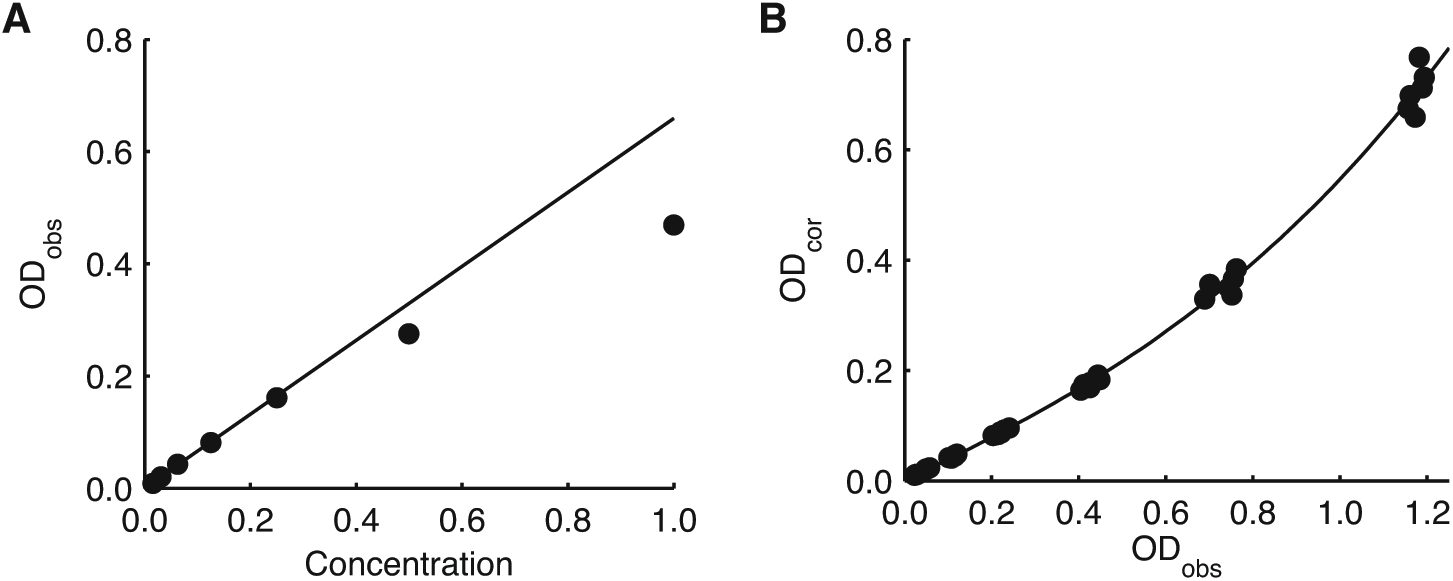
OD_600_ correction for linearity at high cell density. (**A**) OD_600_ measurements after blank subtraction (OD_obs_) from a sample dilution series. Values judged to be in a linear range were fit with a straight line to generate a corrected OD (OD_cor_) for each concentration. (b) ODobs and ODcor values for six dilution series from three independent experiments were fit with a cubic polynomial (OD_cor_ = 0.319 × OD_obs_^3^ + 0.089 × OD_obs_^2^+ 0.959 × ODobs).

**Figure S2.**
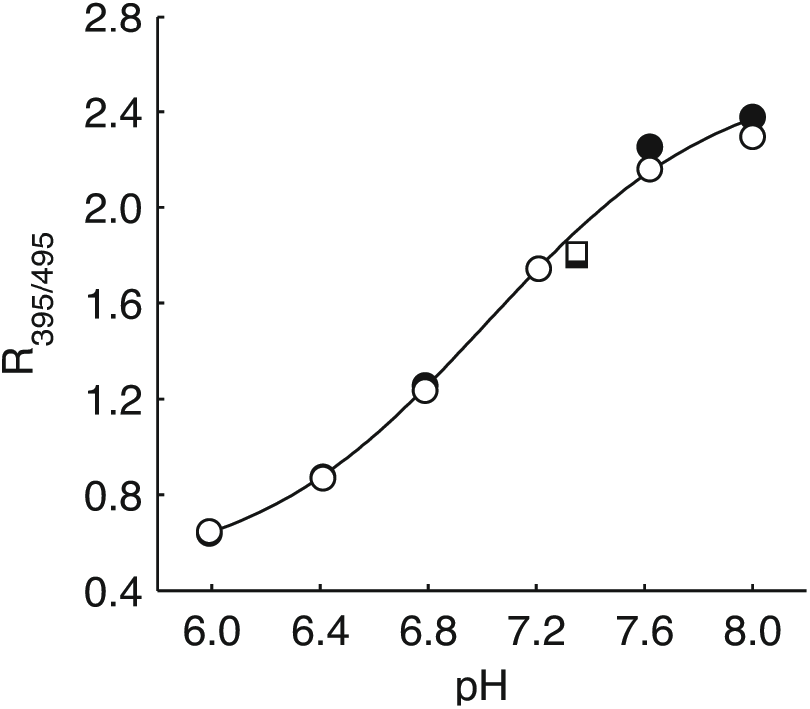
Calibration of pHluorin fluorescence. Cells expressing pHluorin (open symbols) or pHluorin and Lyp1 (closed symbols) were semi-permeabilized using digitonin and diluted into 100 mM KPi (circles) or PBS (squares) buffers. Emission intensity at 508 nm was measured using excitation at 395 and 475 nm, and the ratio (R395M95) plotted against buffer pH. The KPi data were fit with a modified Henderson-Hasselbalch equation: pH = p*K*_a_’ + log_10_((R395/495 – R_min_)/(R_max_ - R_395/495_)), where p*K*a’= −log_10_ of the apparent acid dissociation constant, and R_min_ and R_max_= the R_395/495_ at extreme low and high pH, respectively.

**Figure S3.**
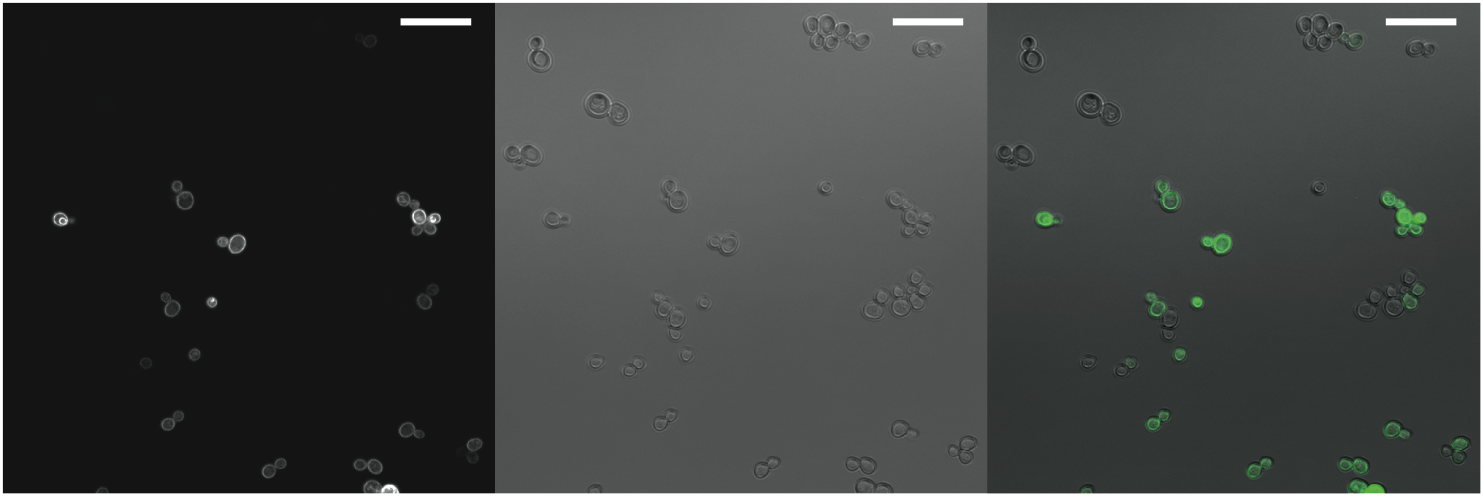
Confocal microscopy images showing the variation in fluorescent signal between cells in the same culture. The strain pictured is expressing Tat2YPet, although similar results were seen for all the overexpression strains. Left = fluorescence, center = brightfield, right = overlay.

**Figure S4.**
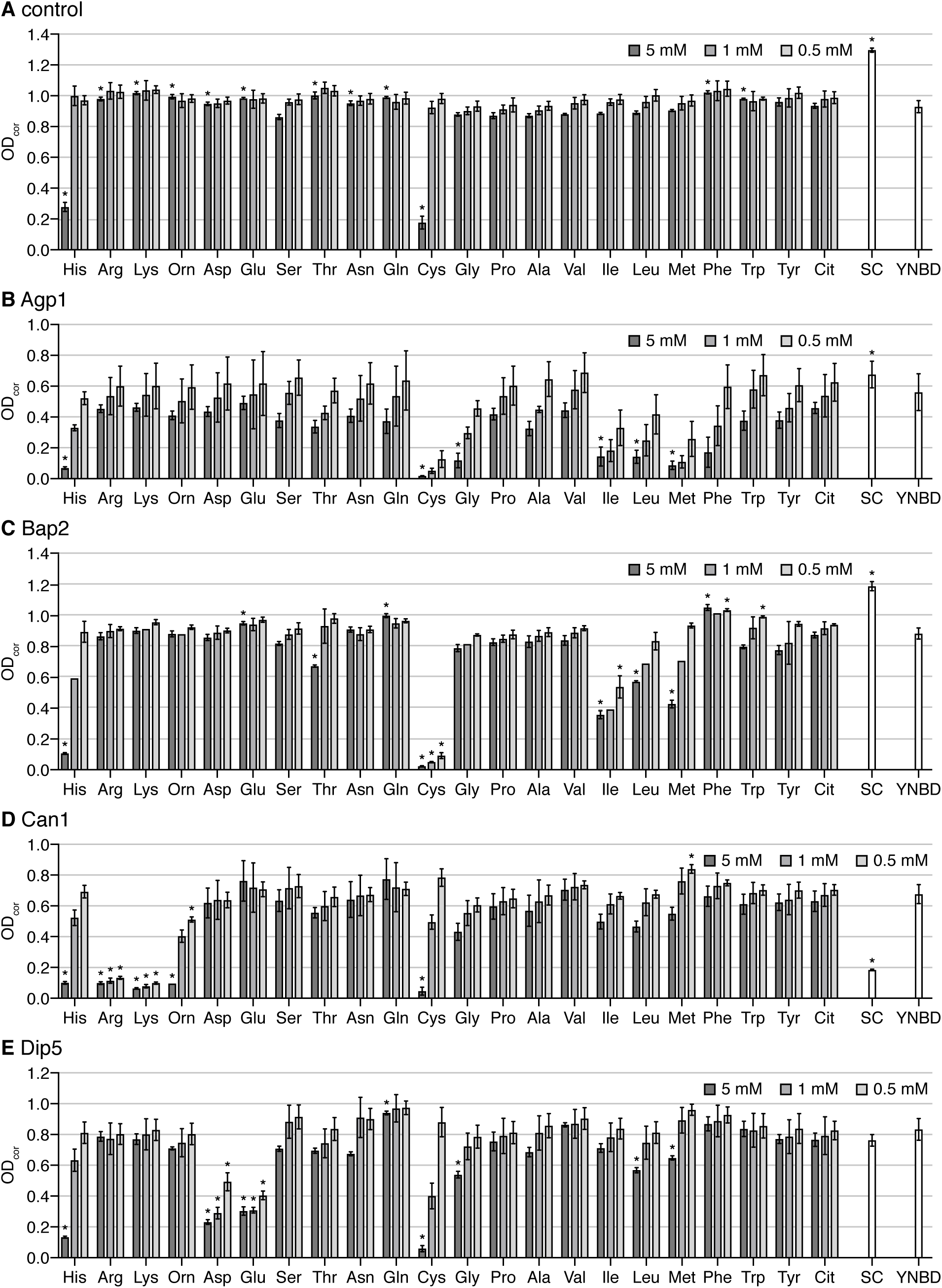
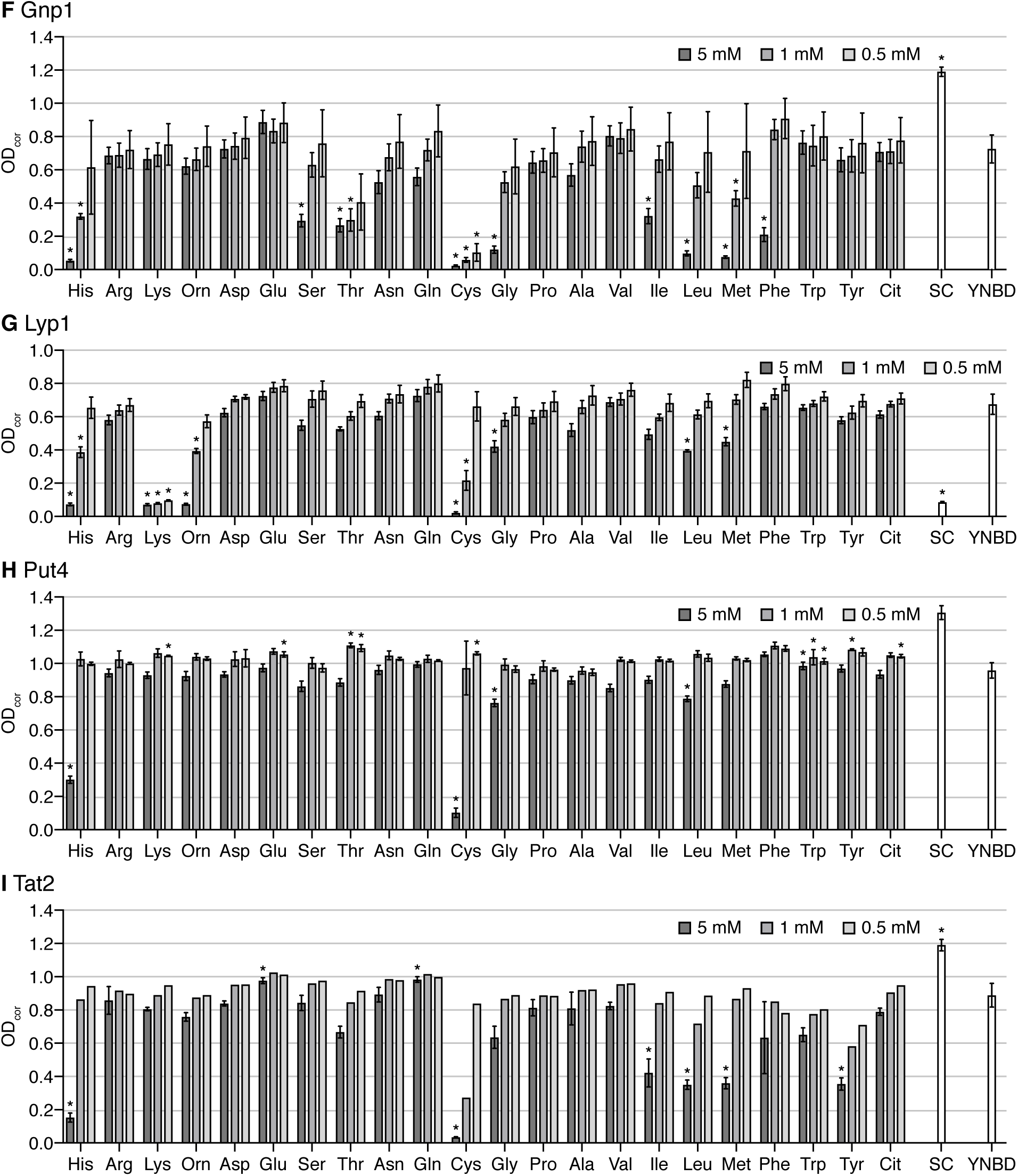
Raw data from experiments presented in Figure 2, showing the effect of amino acids on the growth of strains overexpressing amino acid transporters. BY4709 carrying the empty plasmid pRSII426 was used as a control. All proteins were expressed from the constitutive ADH1 promoter with a C-terminal YPet tag. The proteinogenic amino acids are represented by their three letter codes (see Table 4). Orn, ornithine; Cit, citrulline; SC, synthetic complete media minus uracil; YNBD, minimal medium without amino acids. Amino acids were at final concentrations of 0.5, 1, or 5 mM. OD_cor_ is the measured OD_600_ after 24 h of growth, after blank subtraction and correction for linearity at high cell densities (see Materials and methods). Values shown are the mean ± standard deviation (n = 3 for amino acids and SC, n = 9 for YPD). Due to technical error some results had to be discarded and therefore n = 2 for Bap2 1 mM Gly/His/Ile/Leu/Lys/Met/Orn/Phe, and also for all Tat2 1 mM and 0.5 mM conditions (no error bars are given for these). Although this figure shows the mean of all replicates, the normalized heatmaps in Figure 2 and the statistical analyses were generated by comparing specific pairs of growth values (e.g. YNBD vs YNBD + Cit for samples from the same microplate, inoculated from the same preculture). Asterisks indicate *p*< 0.05 when compared to growth in YNBD (*t*-test).

**Figure S5.**
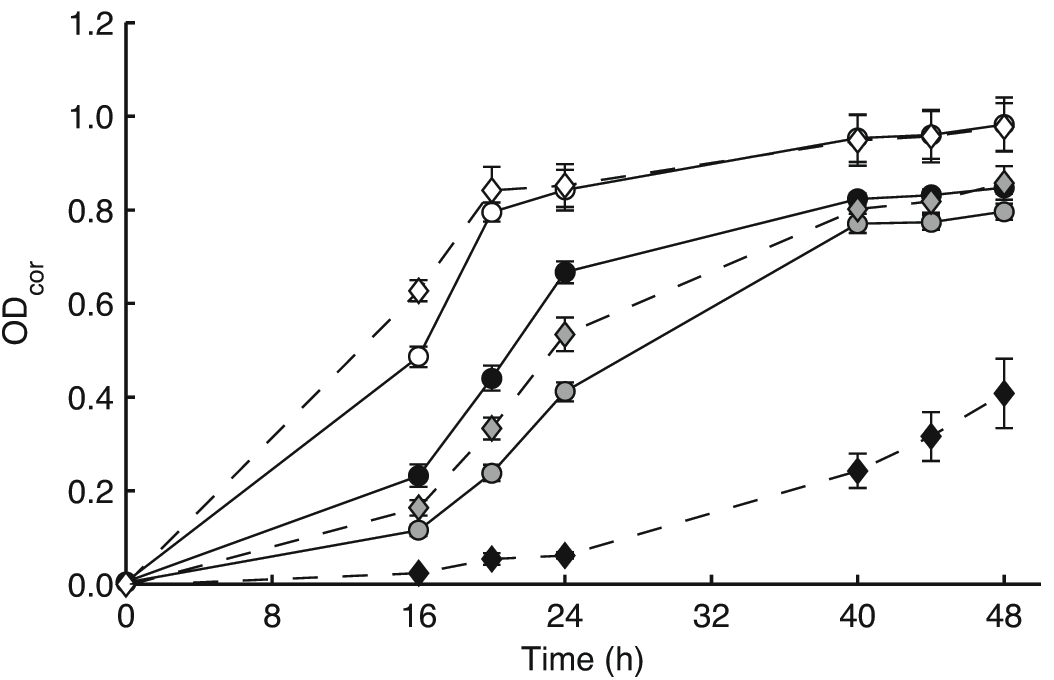
Growth curves from the same experiment shown in Figure 4. Shown here are cultures grown with no external amino acids (circles, solid lines) or with 500 μM arginine (diamonds, dashed lines). Strains without the arginine sensitivity phenotype experience a growth advantage in the presence of external arginine. At t = 24 h, the time-point shown in Figure 4, this advantage is only apparent for the slower growing Lyp1 overexpression strain. Open symbols = BY4709 carrying the empty plasmid pRSII426, closed symbols = cells overexpressing Can1, shaded symbols = cells overexpressing Lyp1. All values shown are the mean of biological triplicates. Error bars represent standard deviation and in some cases are obscured by the data point.

**Figure S6.**
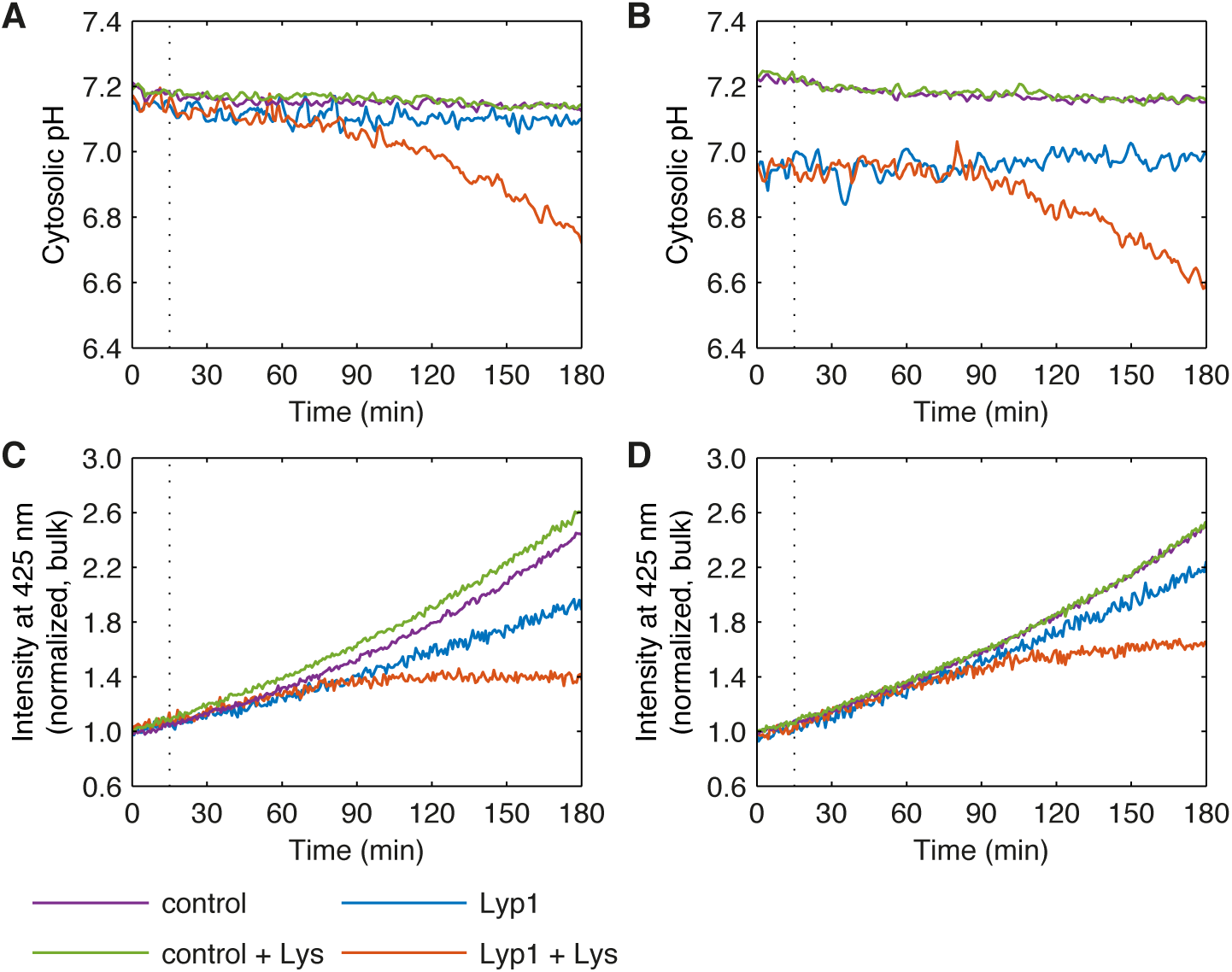
Data from the two experiments used to generate Figure 5. Data in (A) & (C) and (B) & (D) were collected on two separate days, using independent cultures. The pH values were calculated from a single calibration curve shown in Figure S5.

## References

Andréasson C, Neve EPA, Ljungdahl PO. 2004. Four permeases import proline and the toxic proline analogue azetidine-2-carboxylate into yeast. Yeast 21:193–199.

Bender DA. 2012. Amino Acid Metabolism. 3rd ed. John Wiley & Sons, UK.

Bhatnagar RK, Berry A, Hendry AT et al. 1989. The biochemical basis for growth inhibition by L-phenylalanine in *Neisseria gonorrhoeae*. Mol Microbiol 3:429–435.

Bianchi F, Klooster JS, Ruiz SJ et al. 2016. Asymmetry in inward- and outward-affinity constant of transport explain unidirectional lysine flux in *Saccharomyces cerevisiae*. Sci Rep 6:31443.

Bitinaite J, Nichols NM. 2009. DNA cloning and engineering by uracil excision. Curr Protoc Mol Biol Chapter 3: Unit 3.21.

Blau N, van Spronsen FJ, Levy HL. 2010. Phenylketonuria. Lancet 376:1417–1427.

Bonner CA, Jensen RA. 1997. Recognition of specific patterns of amino acid inhibition of growth in higher plants, uncomplicated by glutamine-reversible ‘general amino acid inhibition’. Plant Science 130:133–143.

Bonner CA, Rodrigues AM, Miller JA et al. 1992. Amino acids are general growth inhibitors of *Nicotiana silvestris* in tissue culture. Physiol Plant 84:319–328.

Bonner CA, Williams DS, Aldrich HC et al. 1996. Antagonism by L-glutamine of toxicity and growth inhibition caused by other amino acids in suspension cultures of *Nicotiana silvestris*. Plant Science 113:43–58.

Brachmann CB, Davies A, Cost GJ et al. 1998. Designer deletion strains derived from *Saccharomyces cerevisiae* S288C: a useful set of strains and plasmids for PCR-mediated gene disruption and other applications. Yeast 14:115–132.

Chee MK, Haase SB. 2012. New and redesigned pRS plasmids shuttle vectors for genetic manipulation of *Saccharomyces cerevisiae*. G3 (Bethesda) 2:515–526.

Chen EJ, Kaiser CA 2002. Amino acids regulate the intracellular trafficking of the general amino acid permease of *Saccharomyces cerevisiae*. Proc Natl Acad Sci USA 99:14837–14842.

Christianson TW, Sikorski RS, Dante M et al. 1992. Multifunctional yeast high-copy-number shuttle vectors. Gene 110:119–122.

Cormerais Y, Giuliano S, LeFloch R et al. 2016. Genetic disruption of the multifunctional CD98/LAT1 complex demonstrates the key role of essential amino acid transport in the control of mTORC1 and tumor growth. Cancer Res 76:4481–4492.

Courchesne WE, Magasanik B. 1983. Ammonia regulation of amino acid permeases in *Saccharomyces cerevisiae*. Mol Cell Biol 3:672–683.

Dechant R, Binda M, Lee SS et al. 2010. Cytosolic pH is a second messenger for glucose and regulates the PKA pathway through V-ATPase. EMBO J 29:2515–2526.

Dechant R, Saad S, Ibáñez AJ et al. 2014. Cytosolic pH regulates cell growth through distinct GTPases, Arf1 and Gtr1, to promote Ras/PKA and TORC1 activity. Mol Cell 55:409–421.

Duan Y, Li F, Tan K et al. 2015. Key mediators of intracellular amino acids signaling to mTORC1 activation. Amino Acids 47:857–867.

Durán RV, Oppliger W, Robitaille AM et al. 2012. Glutaminolysis activates Rag-mTORC1 signaling. Mol Cell 47:349–358.

Düring-Olsen L, Regenberg B, Gjermansen C et al. 1999. Cysteine uptake by *Saccharomyces cerevisiae* is accomplished by multiple permeases. Curr Genet 35:609–617.

Englesberg E, Bass R, Heiser W. 1976. Inhibition of the growth of mammalian cells in cuture by amino acids and the isolation and characterization of L-phenylalanine transport. Somatic Cell Genet 2:411–428.

Finley D, Ulrich HD, Sommer T et al. 2012. The ubiquitin-proteasome system of *Saccharomyces cerevisiae*. Genetics 192:319–360.

Ghaddar K, Krammer E-M, Mihajlovic N et al. 2014a. Converting the yeast arginine Can1 permease to a lysine permease. J Biol Chem 289:7232–7346.

Ghaddar K, Merhi A, Saliba E et al. 2014b. Substrate-induced ubiquitylation and endocytosis of yeast amino acid permeases. Mol Cell Biol 34:4447–4463.

Gournas C, Saliba E, Krammer E-M et al. 2016. Function and regulation of fungal amino acid transporters: insights from predicted structure. In Yeast Membrane Transport. Advances in Experimental Medicine and Biology, vol 892, Ramos J, Sychrová H, Kschischo M. (eds). Springer: Cham; 69–106.

Grauslund M, Didion T, Kielland-Brandt MC et al. 1995. *BAP2,* a gene encoding a permease for branched-chain amino acids in *Saccharomyces cerevisiae*. Biochim Biophys Acta 1269:275–280.

Grenson M, Hou C, Crabeel M. 1970. Multiplicity of the amino acid permeases in *Saccharomyces cerevisiae*. IV. Evidence for a general amino acid permease. J Bacteriol 103:770–777.

Grenson M, Mousset M, Wiame JM et al. 1966. Multiplicity of the amino acid permeases in *Saccharomyces cerevisiae*. I. Evidence for a specific arginine-transporting system. Biochim Biophys Acta 127:325–338.

Grenson M. 1966. Multiplicity of the amino acid permeases in *Saccharomyces cerevisiae.* II. Evidence for a specific lysine-transporting system. Biochim Biophys Acta 127:339–346.

Hatakeyama R, Kamiya M, Takahara T et al. 2010. Endocytosis of the aspartic acid/glutamic acid transporter Dip5 is triggered by substrate-dependent recruitment of the Rsp5 ubiquitin ligase via the arrestin-like protein Aly2. Mol Cell Biol 30:5598–5607.

Hinnebusch AG. 2005. Translational regulation of *GCN4* and the general amino acid control of yeast. Annu Rev Microbiol 59:407–450.

Iraqui I, Vissers S, Bernard F et al. 1999. Amino acid signaling in *Saccharomyces cerevisiae:* a permease-like sensor of external amino acids and F-Box protein Grr1p are required for transcriptional induction of the *AGP1* gene, which encodes a broad-specificity amino acid permease. Mol Cell Biol 19:989–1001.

Jauniaux JC, Vandenbol M, Vissers S et al. 1987. Nitrogen catabolite regulation of proline permease in *Saccharomyces cerevisiae*. Cloning of the *PUT4* gene and study of PUT4 RNA levels in wild-type and mutant strains. Eur J Biochem 164:601–606.

Jensen RA, Stenmark-Cox S, Ingram LO. 1974. Mis-regulation of 3-deoxy-D-arabino-heptulosonate 7-phosphate synthetase does not account for growth inhibition by phenylalanine in *Agmenellum quadruplicatum*. J Bacteriol 120:1124–1132.

Jewell JL, Kim YC, Russell RC et al. 2015. Differential regulation of mTORC1 by leucine and glutamine. Science 347:194–198.

Johnson CL, Vishniac W. 1970. Growth inhibition in *Thiobacillus neapolitanus* by histidine, methionine, phenylalanine, and threonine. J Bacteriol 104:1145–1150.

Kanda N, Abe F. 2013. Structural and functional implications of the yeast high-affinity tryptophan permease Tat2. Biochemistry 52:4296–4307.

Kaur J, Bachhawat AK. 2007. Yct1p, a novel, high-affinity, cysteine-specific transporter from the yeast *Saccharomyces cerevisiae*. Genetics 176:877–890.

Keener, J.M, Babst, M. 2013. Quality control and substrate-dependent downregulation of the nutrient transporter Fur4. Traffic 14:412–427.

Krall AS, Xu S, Graeber TG et al. 2016. Asparagine promotes cancer cell proliferation through use as an amino acid exchange factor. Nat Commun 7:11457.

Laplante M, Sabatini DM. 2012. mTOR signaling in growth control and disease. Cell 149:274–293.

Lasko PF, Brandriss MC. 1981. Proline transport in *Saccharomyces cerevisiae*. J Bacteriol 148:241–247.

Leavitt RI, Umbarger HE. 1962. Isoleucine and valine metabolism in *Escherichia coli.* XI. Valine inhibition of the growth of *Escherichia coli* strain K-12. J Bacteriol 83:624–630.

Lin CH, MacGurn JA, Chu T et al. 2008. Arrestin-related ubiquitin-ligase adaptors regulate endocytosis and protein turnover at the cell surface. Cell 135:714–725.

Ljungdahl PO, Daignan-Fornier B. 2012. Regulation of amino acid, nucleotide, and phosphate metabolism in *Saccharomyces cerevisiae*. Genetics 190:885–929.

Ljungdahl PO. 2009. Amino-acid-induced signalling via the SPS-sensing pathway in yeast. Biochem Soc Trans 37:242–247.

MacGurn JA, Hsu P-C, Emr SD. 2012. Ubiquitin and membrane protein turnover: from cradle to grave. Annu Rev Biochem 81:231–259.

Magasanik B, Kaiser CA. 2002. Nitrogen regulation in *Saccharomyces cerevisiae*. Gene 290:1–18.

Melnykov AV. 2016. New mechanisms that regulate *Saccharomyces cerevisiae* short peptide transporter achieve balanced intracellular amino acid concentrations. Yeast 33:21–31.

MiesenbÖck G, De Angelis DA, Rothman JE. 1998. Visualizing secretion and synaptic transmission with pH-sensitive green fluorescent proteins. Nature 394:192–195.

Miles DO, Dyer JK, Wong JC. 1976. Influence of amino acids on the growth of *Bacteroides melaninogenicus*. J Bacteriol 127:899–903.

Mumberg D, Müller R, Funk M. 1995. Yeast vectors for the controlled expression of heterologous proteins in different genetic backgrounds. Gene 156:119–122.

Nguyen AW, Daugherty PS. 2005. Evolutionary optimization of fluorescent proteins for intracellular FRET. Nat Biotechnol 23:355–360.

Nikko E, Pelham HRB. 2009. Arrestin-mediated endocytosis of yeast plasma membrane transporters. Traffic 10:1856–1867.

Nishiuch Y, Sasaki M, Nakayasu M et al. 1976. Cytotoxicity of cysteine in culture media. In Vitro 12:635–638.

Nørholm MHH. 2010. A mutant Pfu DNA polymerase designed for advanced uracil-excision DNA engineering. BMC Biotechnol 10:21–27.

O’Donnell AF, Apffel A, Gardner RG et al. 2010. α-Arrestins Aly1 and Aly2 regulate intracellular trafficking in response to nutrient signaling. Mol Biol Cell 21:3552–3566.

Ohsumi Y, Anraku Y. 1981. Active transport of basic amino acids driven by a proton motive force in vacuolar membrane vesicles of *Saccharomyces cerevisiae*. J Biol Chem 256:2079–2082.

Orij R, Postmus J, Beek Ter A et al. 2009. *In vivo* measurement of cytosolic and mitochondrial pH using a pH-sensitive GFP derivative in *Saccharomyces cerevisiae* reveals a relation between intracellular pH and growth. Microbiology 155:268–278.

Orij R, Urbanus ML, Vizeacoumar FJ et al. 2012. Genome-wide analysis of intracellular pH reveals quantitative control of cell division rate by pHc in *Saccharomyces cerevisiae*. Genome Biol 13:R80.

Österberg M, Kim H, Warringer J et al. 2006. Phenotypic effects of membrane protein overexpression in *Saccharomyces cerevisiae*. Proc Natl Acad Sci USA 103:11148–11153.

Park Y-Y, Sohn BH, Johnson RL et al. 2016. Yes-associated protein 1 and transcriptional coactivator with PDZ-binding motif activate the mammalian target of rapamycin complex 1 pathway by regulating amino acid transporters in hepatocellular carcinoma. Hepatology 63:159–172.

Pickart CM, Rose IA. 1985. Functional heterogeneity of ubiquitin carrier proteins. J Biol Chem 260:1573–1581.

Powis K, De Virgilio C. 2016. Conserved regulators of Rag GTPases orchestrate amino acid-dependent TORC1 signaling. Cell Discov 2:15049.

Regenberg B, Düring-Olsen L, Kielland-Brandt MC et al. 1999. Substrate specificity and gene expression of the amino-acid permeases in *Saccharomyces cerevisiae*. Curr Genet 36:317–328.

Regenberg B, Holmberg S, Olsen LD et al. 1998. Dip5p mediates high-affinity and high-capacity transport of L-glutamate and L-aspartate in *Saccharomyces cerevisiae*. Curr Genet 33:171–177.

Risinger AL, Cain NE, Chen EJ et al. 2006. Activity-dependent reversible inactivation of the general amino acid permease. Mol Biol Cell 17:4411–4419.

Ro D-K, Ouellet M, Paradise EM et al. 2008. Induction of multiple pleiotropic drug resistance genes in yeast engineered to produce an increased level of anti-malarial drug precursor, artemisinic acid. BMC Biotechnol 8:83.

Rowley D. 1953a. Inhibition of *E. coli* strains by amino-acids. Nature 171:80–81.

Rowley D. 1953b. Interrelationships between amino-acids in the growth of coliform organisms. J Gen Microbiol 9:37–43.

Sanayama Y, Matsumoto A, Shimojo N et al. 2014. Phenylalanine sensitive K562-D cells for the analysis of the biochemical impact of excess amino acid. Sci Rep 4:6941.

Sato T, Ohsumi Y, Anraku Y. 1984. Substrate specificities of active transport systems for amino acids in vacuolar-membrane vesicles of *Saccharomyces cerevisiae.* Evidence of seven independent proton/amino acid antiport systems. J Biol Chem 259:11505–11508.

Saudubray J-M, Baumgartner MR, Walter J eds. 2016. Inborn Metabolic Diseases: Diagnosis and Treatment. 6 ed. Springer Berlin, Heidelberg, Germany.

Sáenz DA, Chianelli MS, Stella CA. 2014. L-Phenylalanine transport in *Saccharomyces cerevisiae:* Participation of *GAP1, BAP2,* and *AGP1*. J Amino Acids 2014:283962.

Schmidt A, Hall MN, Koller A. 1994. Two FK506 resistance-conferring genes in *Saccharomyces cerevisiae, TAT1* and *TAT2,* encode amino acid permeases mediating tyrosine and tryptophan uptake. Mol Cell Biol 14:6597–6606.

Schreve J, Garrett JM. 1997. The branched-chain amino acid permease gene of *Saccharomyces cerevisiae, BAP2,* encodes the high-affinity leucine permease (S1). Yeast 13:435–439.

Schreve JL, Sin JK, Garrett JM. 1998. The *Saccharomyces cerevisiae YCC5* (YCL025c) gene encodes an amino acid permease, Agp1, which transports asparagine and glutamine. J Bacteriol 180:2556–2559.

Sherman F. 2002. Getting started with yeast. Meth Enzymol 350:3–41.

Stoll KE, Brzovic PS, Davis TN et al. 2011. The essential Ubc4/Ubc5 function in yeast is HECT E3-dependent, and RING E3-dependent pathways require only monoubiquitin transfer by Ubc4. J Biol Chem 286:15165–15170.

Stracka D, Jozefczuk S, Rudroff F et al. 2014. Nitrogen source activates TOR (target of rapamycin) complex 1 via glutamine and independently of Gtr/Rag proteins. J Biol Chem 289:25010–25020.

Sumrada R, Cooper T. 1976. Basic amino acid inhibition of growth in *Saccharomyces cerevisiae*. Biochem Biophys Res Commun 68:598–602.

Sychrová H, Chevallier MR. 1993. Cloning and sequencing of the *Saccharomyces cerevisiae* gene *LYP1* coding for a lysine-specific permease. Yeast 9:771–782.

Sychrová H, Matějčková A, Kotyk A. 1993. Kinetic properties of yeast lysine permeases coded by genes on multi-copy vectors. FEMS Microbiol Lett 113:57–61.

Tarsio M, Zheng H, Smardon AM et al. 2011. Consequences of loss of Vph1 protein-containing vacuolar ATPases (V-ATPases) for overall cellular pH homeostasis. J Biol Chem 286:28089–28096.

Usami Y, Uemura S, Mochizuki T et al. 2014. Functional mapping and implications of substrate specificity of the yeast high-affinity leucine permease Bap2. Biochim Biophys Acta 1838:1719–1729.

Vandenbol M, Jauniaux JC, Grenson M. 1989. Nucleotide sequence of the *Saccharomyces cerevisiae PUT4* proline-permease-encoding gene: similarities between CAN1, HIP1 and PUT4 permeases. Gene 83:153–159.

Warringer J, Blomberg A. 2003. Automated screening in environmental arrays allows analysis of quantitative phenotypic profiles in *Saccharomyces cerevisiae*. Yeast 20:53–67.

Watanabe D, Kikushima R, Aitoku M et al. 2014. Exogenous addition of histidine reduces copper availability in the yeast *Saccharomyces cerevisiae*. Microbial Cell 1:241–246.

Wenzel DM, Lissounov A, Brzovic PS et al. 2011. UBCH7 reactivity profile reveals parkin and HHARI to be RING/HECT hybrids. Nature 474:105–108.

White MA, Clark KM, Grayhack EJ et al. 2007. Characteristics affecting expression and solubilization of yeast membrane proteins. J Mol Biol 365:621–636.

Wilson DM, Kusch M, Flagg-Newton JL et al. 1980. Control of lactose transport in *Escherichia coli*. FEBS Letters 117 Suppl:K37–44.

Wilson DM, Putzrath RM, Wilson TH. 1981. Inhibition of growth of *Escherichia coli* by lactose and other galactosides. Biochim Biophys Acta 649:377–384.

Zheng L, Zhang W, Zhou Y et al. 2016. Recent advances in understanding amino acid sensing mechanisms that regulate mTORC1. Int J Mol Sci 17:1636.

Zhu X, Garrett J, Schreve J et al. 1996. GNP1, the high-affinity glutamine permease of *S. cerevisiae*. Curr Genet 30:107–114.

